# EEG Biomarkers of Age-Related Memory Change

**DOI:** 10.1101/2024.08.20.608804

**Authors:** Adam W. Broitman, M. Karl Healey, Michael J. Kahana

## Abstract

The current study investigates whether electroencephalographic (EEG) activity reflects age-related memory changes during encoding. We recorded scalp EEG in 151 young adults (aged 18-30) and 37 older adults (aged 60-85) as they memorized lists of words. Subjects studied the word lists either under full attention or while performing a secondary task that required them to make semantic judgments about each word. Although the secondary task reduced recall among all subjects, differences in recall performance between the age groups were smaller when participants performed a secondary task at encoding. Older adults also exhibited distinct neural subsequent memory effects, characterized by less negativity in the alpha frequencies compared to young adults. Multivariate classifiers trained on neural features successfully predicted subsequent memory at the trial level in both young and older adults, and captured the differential effects of task demands on memory performance between young and older adults. The findings indicate that neural biomarkers of successful memory vary with both cognitive aging and task demands.

**Public significance:** The current study investigates the EEG spectral biomarkers of memory encoding in young and older adults, and identifies key features of neural activity associated with age-related memory change. We further find that age-related memory differences are smaller when participants perform a semantic judgment task during encoding, and that multivariate classifiers trained on the EEG data predict these effects.

## Introduction

The ability to learn and retain new information declines across the human lifespan (Nilsson, 2003; Salthouse, 2016). A rich literature has aimed to characterize the neurocognitive processes involved in this change using a variety of approaches (Cabeza et al., 2004; Daselaar et al., 2006; Dennis et al., 2007; Kukolja et al., 2016). Spectral analyses of electroencephalogram (EEG) recordings have provided critical insights into the effects of aging on memory processes, linking age-related deficits with specific changes in spectral activity and ERP components (Friedman, 2013; Karrasch et al., 2004; Strunk and Duarte, 2019). However, several key questions about the EEG biomarkers of age-related memory change remain unanswered. Specifically, it is unclear whether the spectral activity that predicts memory encoding in young adults similarly predicts encoding in older adults, and whether any age-related differences in these biomarkers are sensitive to task demands. The current study investigates these questions using multivariate spectral decomposition of EEG in a study of free recall.

### Do EEG biomarkers of memory encoding vary with age and task demands?

Studies aiming to characterize the role of neural variability in memory encoding have identified patterns of brain activity, known as neural *subsequent memory effects* (SMEs), that can help predict whether a subject will remember information about a particular event (Kim, 2011; Long et al., 2014; Paller et al., 1987). For example, several scalp EEG studies with young adults have found that increased activity in the gamma band (30-100 Hz), coupled with reduced activity in the alpha band (8-14 Hz), indicate a greater likelihood of remembering items presented in that moment (Katerman et al., 2022; Broitman and Swallow, 2023; Long et al., 2014; Weidemann and Kahana, 2021). Some evidence suggests that dividing attention across multiple tasks or study items disrupts this activity, and its relationship with memory encoding. Sederberg et al. (2006) reported that in free recall experiments, gamma SMEs weaken in late serial positions as subjects simultaneously process multiple study items from earlier list positions. In a scalp EEG study of free recall, Broitman and Swallow (2023) found that, while gamma activity was associated with later recall under single-task encoding conditions, it did not predict subsequent recall when subjects divided attention across tasks by performing target detection on colored squares during word list encoding. Long and Kahana (2017) also reported that gamma activity at encoding predicted the tendency to temporally cluster word recalls under single task encoding, but not when subjects performed a semantic processing task during word presentation (e.g., “Is this item living/nonliving?”). Together, these studies indicate that the neural predictors of memory encoding in young adults are sensitive to task demands and cognitive load.

Most studies aiming to identify the neural correlates of age-related memory change have primarily examined memory failures that occur during retrieval. EEG applications have typically examined event-related potentials (ERPs), linking age-related memory changes to reductions in components associated with recollection and memory specificity (Friedman, 2013; Finnigan et al., 2011; Li et al., 2004). However, ERPs provide limited resolution in measuring neural activity that is not time-locked to a stimulus presentation (Jacobs et al., 2006; Bastiaansen et al., 2011). Spectral decomposition of EEG can measure this activity, and has been successful in detecting neurocognitive impairment (McBride et al., 2014). The few existing studies that have used spectral decomposition of EEG to examine age-related differences at encoding (Karrasch et al., 2004; Strunk and Duarte, 2019) reported subtle age-related differences in alpha activity. However, these studies relied on old/new recognition memory tests, which typically report minimal age-related impairments and rarely capture age-related memory changes as effectively as the free recall paradigm (Rhodes et al., 2019). With little research available, it is therefore unclear how the spectral EEG predictors of memory encoding vary between young and older adults in a free recall task.

### The Current Study

The current study investigates the neural correlates of memory encoding using data from the Penn Electrophysiology of Encoding and Retrieval Study (PEERS, Kahana et al., 2022), a multi-session experiment with young (age 18-30) and older (age 61-85) adults. Subjects encoded lists of words either on their own (No Task condition), or while performing a secondary semantic judgement task on the words (Task condition), and then vocally recalled as many words from the just-encoded list as they could remember. Our primary questions are: 1) How do EEG spectral biomarkers of encoding success vary across age groups? 2) Do young and older adults demonstrate similar disruptions to encoding when the experiment imposes a semantic processing task? and 3) Can scalp EEG signals capture any potential interactions of cognitive aging and semantic processing?

Prior analysis of the PEERS dataset have already found that young adults recall fewer items when they perform semantic judgments on each word during encoding (Long and Kahana, 2017). In the absence of experimenter-imposed processing tasks, young adults tend to adopt effective order-based encoding strategies. The imposition of a processing task can disrupt such strategies and produce dual-task encoding interference, impairing recall performance (Mundorf et al., 2022). However, it is unclear whether young and older adults demonstrate the same effects of experimenter-imposed semantic processing. Existing literature suggests that older adults are particularly vulnerable to language processing deficits while performing multiple tasks simultaneously (Park et al., 1989; Kemper et al., 2010), suggesting that the secondary task will impair older adults’ memory more than that of the young adults. However, in these previous studies, the secondary tasks were unrelated to the encoding stimuli (e.g., digit monitoring, visual target detection). Results may differ when the secondary task involves performing semantic judgments on the to-be-encoded stimuli, especially given the extensive evidence that semantic processing remains intact among older adults (Verhaeghen, 2003; Mayr and Kliegl, 2000; Lacombe et al., 2015). If there are smaller age-related differences in Task lists than in No Task lists, then the results would support prior assertions that aging accompanies a reduced reliance on order-based strategies for word list encoding (Sanders et al., 1980; Jackson and Schneider, 1982; Healey and Kahana, 2016).

Spectral analysis of scalp EEG may provide critical insights into the effects of aging on encoding processes. Some theoretical frameworks account for age-related memory change by positing that aging reduces the ability to inhibit the processing of task-irrelevant information, thereby increasing cognitive load during encoding (Craik and Byrd, 1982; Hasher and Zacks, 1988). If this is the case, then we might expect the neural biomarkers of memory encoding for older adults on the No Task lists to resemble those of the young adults during Task lists. In young adults, increasing task demands during encoding can weaken the relationship between gamma activity and mnemonic success (Broitman and Swallow, 2023; Long and Kahana, 2017). Therefore, if older adults struggle to sustain attention to the task and to inhibit task-irrelevant information, then they may demonstrate a reduced gamma SME consistent with increased cognitive load. Aging is also associated with decreased activity in both the alpha and gamma bands (Voytek et al., 2015; Babiloni et al., 2006; Murty et al., 2020), which could indicate that activity in these frequencies will more weakly associate with memory in older adults. Older adults with preserved memory performance have demonstrated suppressed alpha activity during word list encoding, possibly reflecting the deployment of compensatory mnemonic strategies (Healey and Kahana, 2020), and further supporting our expectation of age-related disruptions to alpha band SMEs.

Any potential main effects and interactions of age and task demands on memory performance might be too subtle to detect in univariate comparisons of broad, band-averaged spectral power estimates (Li et al., 2024; Kragel et al., 2017). However, multivariate EEG classifiers can predict encoding success at the single-trial level by aggregating thousands of features of neural activity across multiple sessions (Ezzyat et al., 2018; Phan et al., 2019). These tools may offer superior sensitivity in identifying the features that predict encoding in young and older adults, and in capturing the interacting effects of aging and task demands on memory performance. We will therefore apply a combination of univariate and multivariate analysis of the spectral data to optimally capture the neural effects of age and task demands on memory performance.

## Methods

### Transparency and Openness

Data, study materials, and key EEG data analysis scripts from the current study are available for download at [blinded]. The current study was not preregistered.

### Subjects

Subjects were 188 right-handed, native English speaking, neurocognitively healthy adults. 151 of these subjects were 18-30 years old (young adult cohort), and 37 subjects were 61-85 years old (older adult cohort). The mean age among younger subjects was 22.4 (SD = 3.0), and the mean age among older adult subjects was 69.2 (SD = 6.5). Older adults had an average of 17.82 (SD = 2.51) years of education, and young adults had an average of 14.94 (SD = 1.98) years of education. Additional demographic information regarding the subject samples can be found in Healey and Kahana (2016). Subjects were paid between $10-20 per session for their participation in the study, and were offered a $5 bonus for minimizing their eye blinks during the experiment. The University of [blinded] IRB reviewed and approved all recruitment and testing procedures.

### Experimental Materials and Equipment

Subjects sat unconstrained in a normally lit faraday enclosure, approximately 90 cm away from a ViewSonic 17” LCD monitor (1024×768 pixels, 75Hz refresh rate). All experiments were programmed using Python.

Word lists were constructed using words drawn from a pool of 1638 nouns taken from the University of South Florida free-association norms (Nelson et al., 2004). We constructed the lists such that adjacent items would have a variable degree of semantic similarity to one another. Using the Word Association Spaces (WAS; Steyvers et al., 2004), word pairs were segmented into four similarity bins ranging from high (*cosθ > .*8) to low (*cosθ < .*14). Each list included two word pairs chosen from each bin, with words from one pair placed at adjacent serial positions and words from the other pair separated by at least two serial positions (see Kahana et al., 2022).

### Free Recall Task

The data reported in this article were collected as part of the PEERS experiment Kahana et al. (2022). In each of seven sessions, subjects completed 16 cycles of the verbal free recall task. Each trial (duration = 3800-4200 ms) consisted of a word presented at the center of the screen for 3000 ms. Following word presentation, the screen remained blank for a jittered (i.e., variable) interstimulus period of 800-1200 ms. Following the presentation of the final list item, a jittered delay of 1200-1400 ms occurred, followed by an auditory tone and the appearance of a row of asterisks onscreen. At this time, subjects had 75 s to vocally recall as many items as they could remember from the previous word list.

During four out of the 16 word lists, subjects encoded the words on their own, without a secondary task (No Task lists). During the other 12 lists (Task lists), word presentation included a secondary task in which subjects made judgements regarding the size or animacy of each word (i.e., is the item living/nonliving, can the item fit into a shoebox). The font color of the presented word indicated which task participants should perform (e.g., green font = living/nonliving, purple font = shoebox). Subjects provided their responses by pressing a key on a keyboard. For size judgments, subjects pressed the “K” key to indicate large items and the “L” key to indicate small items. For animacy judgments, subjects pressed the “,” key for living items and the “.” key for nonliving items. Of the 12 lists involving a judgement task, four included size judgements only, four included animacy judgements only, and four had the subjects switch between the tasks throughout list encoding. Because rapid task switching may involve additional neurocognitive processes that are beyond the scope of the current investigation, we excluded the switch task lists from our analyses.

### EEG methods

#### Data collection and preprocessing

We collected EEG data using a 129-channel EGI geodisic system at a sampling rate of 500 Hz. During the experiment, subjects attempted to minimize bodily movements and eye blinks. Following data collection, we applied a Butterworth notch filter at 60 Hz to eliminate electrical noise, as well as a low-pass filter at 1 Hz to eliminate slow drift artifacts. We identified bad channels as those with extremely high or low (*|z| >* 3) log-variance, or which had an extremely high (*z >* 3) Hurst exponent relative to other channels. After dropping the bad channels, we re-referenced the recordings to the common average across all remaining electrodes. To remove eyeblinks and other artifacts, we used an automatic artifact detection algorithm based on independent components analysis (Nolan et al., 2010). Additional details regarding artifact detection and rejection in the current dataset are available in Healey and Kahana (2020).

#### Spectral density analyses

For each trial, we computed power spectral density at 46 logarithmically spaced frequencies ranging from 2-100 Hz using a Morlet wavelet transformation with a wave number of 5. We then decimated power values by a factor of 32, resulting in a final sampling rate of 16 Hz. We selected 25 time points between 0 and 1600 ms relative to stimulus onset at each trial, because several previous scalp EEG studies reported SMEs within this window (Long et al., 2014; Broitman and Swallow, 2023; Weidemann and Kahana, 2021; Katerman et al., 2022). We then Z-scored power within each channel and frequency separately for each session using the mean and standard deviation of power values across all trials. Importantly, when comparing differences between age groups and serial positions *within* list conditions, we separately z-scored task and no-task trials. For comparisons *across* list conditions, we jointly z-scored the data from Task and No Task lists.

After computing spectral densities, we grouped power into two frequency bands of interest: alpha (8-14 Hz) and gamma (30-100 Hz). We selected these bands based on prior studies consistently reporting significant SMEs at these frequencies (Long et al., 2014; Broitman and Swallow, 2023; Weidemann and Kahana, 2021). For univariate analyses, we selected eight regions of interest (ROI) based on previous memory studies using similar EEG caps (Fig. 3A; Weidemann et al., 2009; Long et al., 2014; Katerman et al., 2022) We then averaged power over each trial from 0-1600 ms post-word onset across the preselected ROI channels.

To investigate differences in power between age groups (older/young), subsequent memory (recalled/nonrecalled), and list condition (No Task/Task), we constructed several mixed-effects linear models. We included age group, subsequent memory, and list condition as fixed effects. The model included random effects to account for differences in the effect of recall status on power across subjects, and across individual words. When comparing effects within list condition, we constructed the models using the following notation: Power *∼*AgeGroup *×* Recalled *×* List Condition +(1+ Recalled *|* Subject) + (1+ Recalled *|* ItemID)

All models were constructed in R using the lme4 package (Bates et al., 2015), and were further characterized using the anova function.

#### Multivariate logistic analyses

We trained subject-specific L2-penalized logistic regression classifiers on neural features extracted from all Task and No Task trials (as with the univariate analyses, switch task trials were excluded). To generate the data to classify mnemonic success, we sampled time-frequency data from all 46 frequencies and 124 scalp electrodes. We averaged the features over time points within a trial to provide a single value per frequency and electrode at each encoding event. These analyses yielded a total of 5,704 features.

Binary labels indicated whether the word presented at the trial was recalled or nonrecalled. To determine the optimal penalization hyperparameter, we conducted a grid search for inverse regularization parameter, *C*, with 10 logarithmically spaced values from 10*^−^*^8^ to 10*^−^*^1^. This procedure produced a *C* parameter value of 1.67 *×* 10*^−^*^4^, which optimized classifier performance. For each subject, we performed a leave-one-out cross-validation and bootstrapping test on each of the seven sessions to evaluate the classifier’s generalizability across sessions. To assess classifier performance, we computed the area under the receiver operating characteristic curve (AUC) for the predicted probabilities of remembering encoding events in the hold-out sessions.

## Results

### Behavioral Results

We first sought to characterize the effects of performing a secondary task on free recall performance. Accuracy on the semantic judgement tasks was high, with 86% accuracy on size judgements and 88% accuracy on animacy judgments. Healey and Kahana (2016) extensively characterized differences in free recall performance between older and young adult subjects in this experiment. However, those analyses compared free recall performance between young and older adults using data from all lists, including those that incorporated a secondary encoding task. Figure 2 compares recall performance across age groups and list conditions. To test the effects of age group and list condition on recall performance, we constructed a logistic linear mixed effects model with recall status (recalled/unrecalled) as a binomial dependent variable, age group and list type as fixed effects, and a random effect of list condition nested within subjects. Joint tests revealed that recall performance was lower among older adults, *F* = 78*, p < .*0001. As Long and Kahana (2017) previously reported, overall recall was lower in Task lists than in No Task ones, *F* = 109*, p < .*0001. Surprisingly however, differences in recall performance between the age groups were smaller on Task lists than on No Task lists, age *×* list condition interaction, *F* = 4.6*, p* = .03. Differences in recall performance between the age groups were therefore smaller on Task lists than on No Task lists.

**Figure 1.**
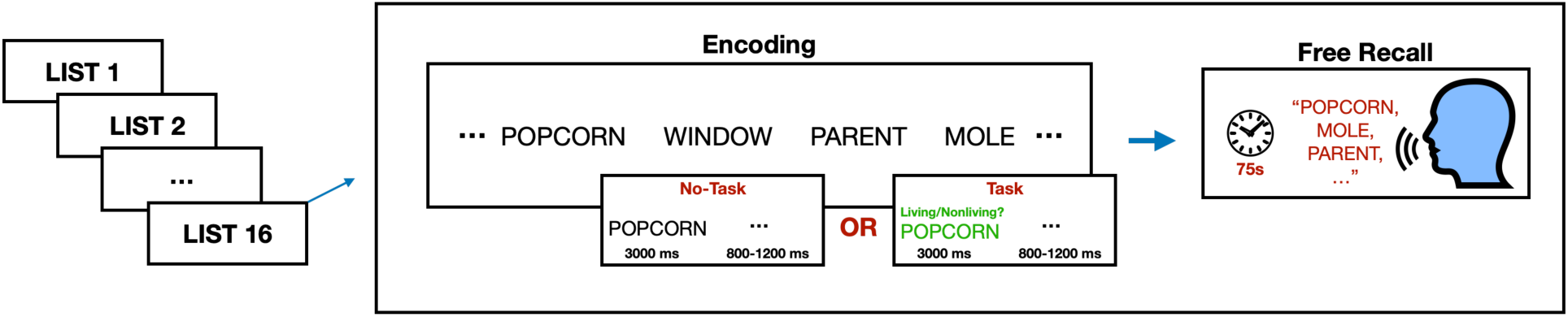
Overview of Experimental Paradigm. Subjects performed free recall on lists of 16 words. Subjects either encoded the words on their own, without a secondary task (No Task lists), or while performing a secondary judgement task on the meaning of the words (Task lists). Font color of the presented words indicated which judgment task participants were to perform.

**Figure 2.**
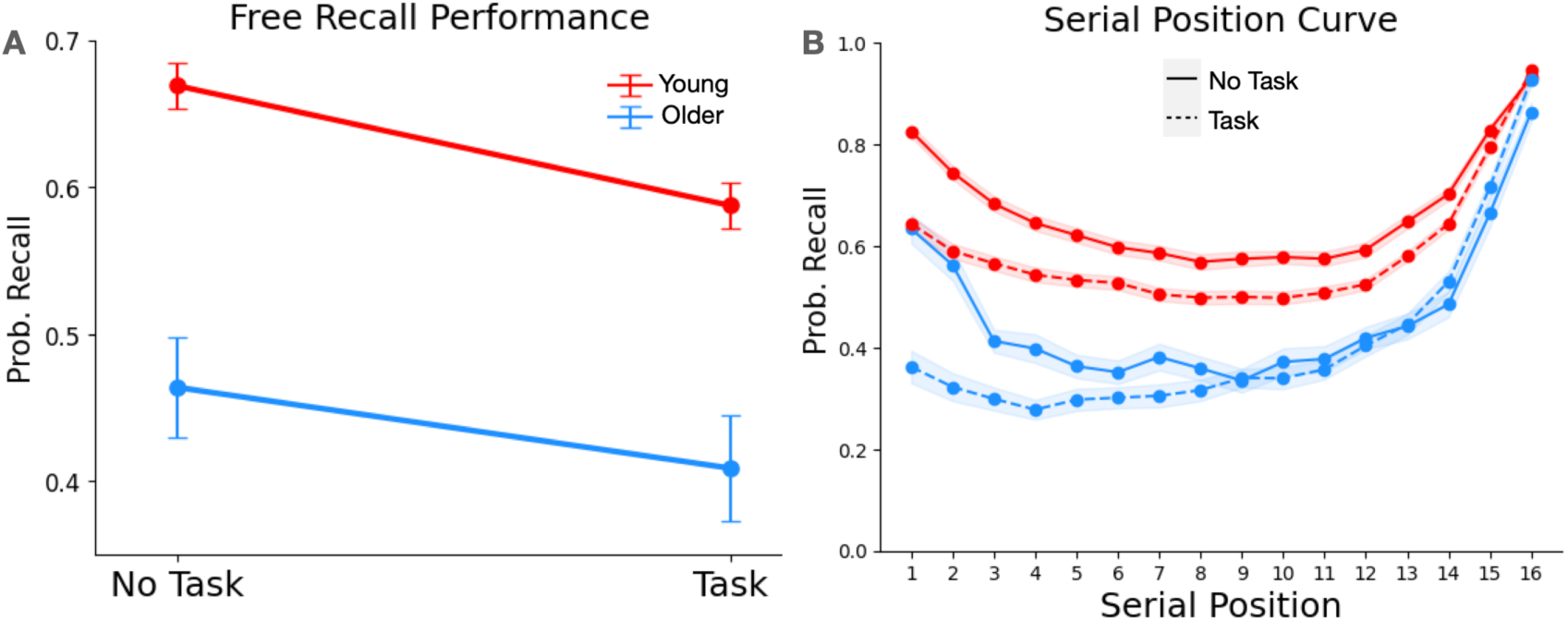
Recall performance across age groups and list conditions. **A.** Average recall performance in Task and No Task lists, plotted separately for young adults (red) and older adults (blue). **B.** Serial position function by age group and list condition. Error bands represent one standard error of the mean.

### EEG Results

#### Neural SMEs across age groups

Figure 3A presents the subsequent memory effects for young and older subjects. We sought to evaluate whether EEG activity varied across trials that were subsequently recalled vs. nonrecalled, and whether these effects varied across age groups. To create this figure, for each subject we computed two-samples *t*-tests between recalled and nonrecalled trials at each frequency and ROI. We then performed a one-sample *t*-test of all subjects’ *t* values to obtain an across-subject *t*-score, which we plot in the spectrograms. Because prior scalp EEG studies have reported SMEs in the gamma and alpha bands (Long et al., 2014; Broitman and Swallow, 2023; Weidemann and Kahana, 2021), our mixed-effects linear regression analyses focused on these frequencies. We did not observe an overall SME in the gamma band, *F < .*1*, p* = .79. We did, however, observe a significant, negative alpha SME, *F* (1, 211130) = 18.8*, p < .*001. This SME was significantly more negative among young adults than among older ones, age *×* recall status interaction, *F* (1, 195) = 3.9*, p < .*05. These results suggest that the relationship between alpha activity and memory encoding may weaken with age.

**Figure 3.**
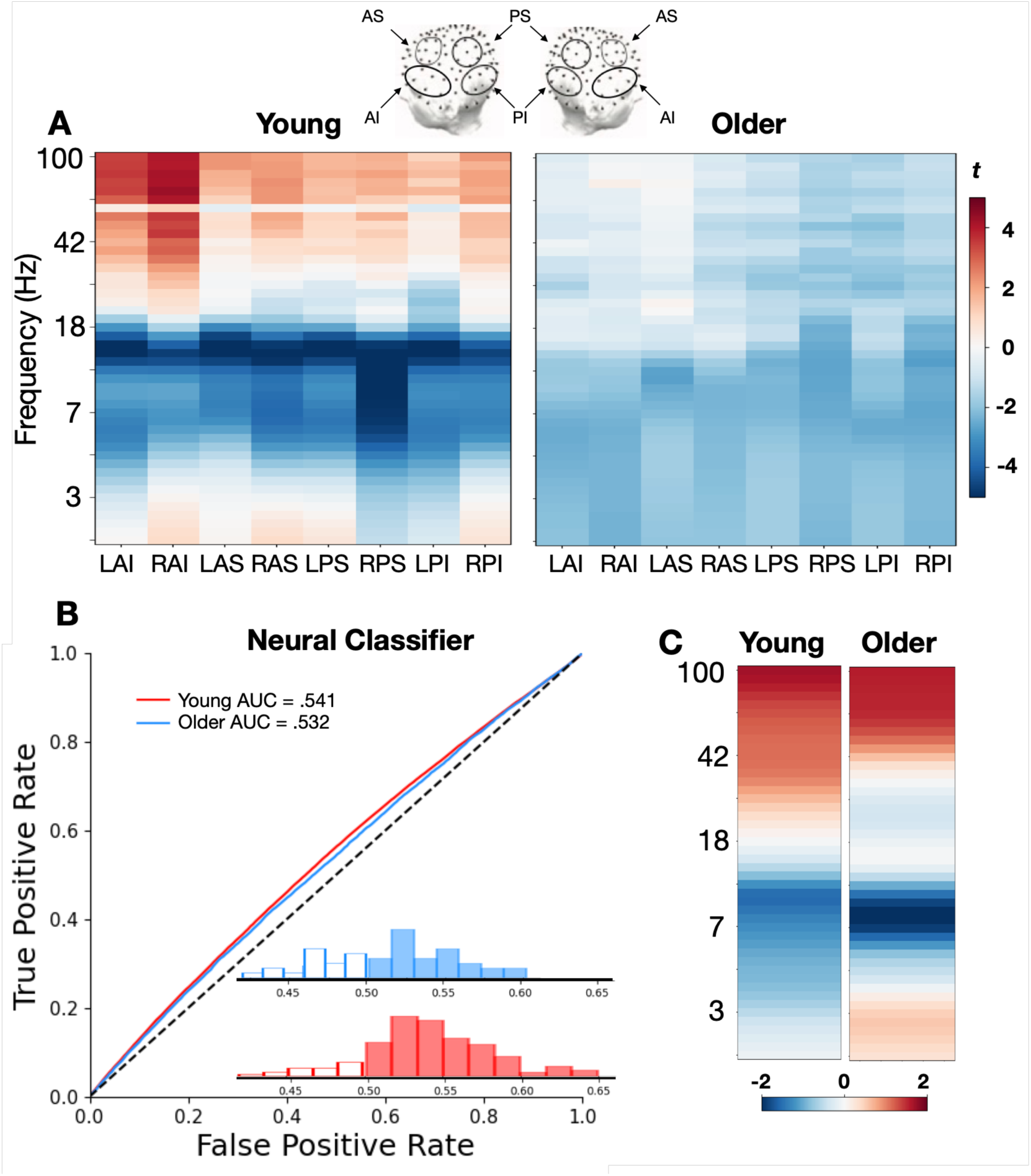
Spectral subsequent memory effects. A) Across-subject t-statistic for recalled - nonrecalled z-power at each computed frequency and. ROI include all permutations of Left/Right (L/R), Anterior/Posterior (A/P), and Superior/Inferior (S/I) SMEs are shown separately for young (left) and older (right) subjects across encoding trials. B) ROC curves showing classifier performance in young (red) and older (blue) adults, as well as histograms of AUCs within each age group. C). Classifier activations constructed using a forward model based on the learned weights (Haufe et al., 2014).

We were additionally interested in whether neural classifiers could predict encoding success with equal accuracy among young and older adults. The classifiers predicted encoding success significantly above chance levels across both age groups, with an average AUC of .54, *t*(178) = 15*, p < .*0001 (Fig. 3B). Figure 3C presents activations constructed from a forward model based on the learned weights of the individual classifiers across all scalp electrodes (Haufe et al., 2014). Despite observable differences in the weighting of frequencies for memory classification, an independent samples *t*-test indicated that goodness-of-fit did not significantly vary across the age groups, *t*(71) = 1.4*, p* = .15, suggesting that neural decoders can predict older and young adults’ memory performance with similar accuracy.

#### Neural SMEs across list conditions

If spectral SMEs vary across age groups, do they vary across list conditions as well, and do any potential effects of list type interact with age? Overall levels of both alpha and gamma activity increased in Task lists relative to No Task lists, *F ^′^s >* 195*, p^′^s < .*0001. This increase in alpha power across list types was larger among older adults as seen in a reliable interaction of age and list type, *F* (1, 211073) = 17.23*, p < .*0001.

The lack of an observed SME in the gamma frequencies may be due to the differential relationship between gamma power and subsequent memory across list conditions. Consistent with this possibility, an interaction of recall status and list condition indicated that gamma predicts subsequent recall positively in the No Task lists, but negatively within Task lists, *F* (1, 205720) = 21.1*, p < .*0001 (Fig. 4). To investigate whether any significant gamma band SMEs were present within list conditions, we separately modeled gamma power within Task and No Task lists. Joint tests on the mixed linear models revealed a significantly positive gamma SME in the No Task lists, *F* (1, 178) = 5.6*, p < .*05, and a significantly negative gamma SME in the Task lists, *F* (1, 189) = 6.7*, p* = .01. This interaction is consistent with existing evidence that the relationship between gamma power and memory encoding changes under dual-task conditions (Broitman and Swallow, 2023).

**Figure 4.**
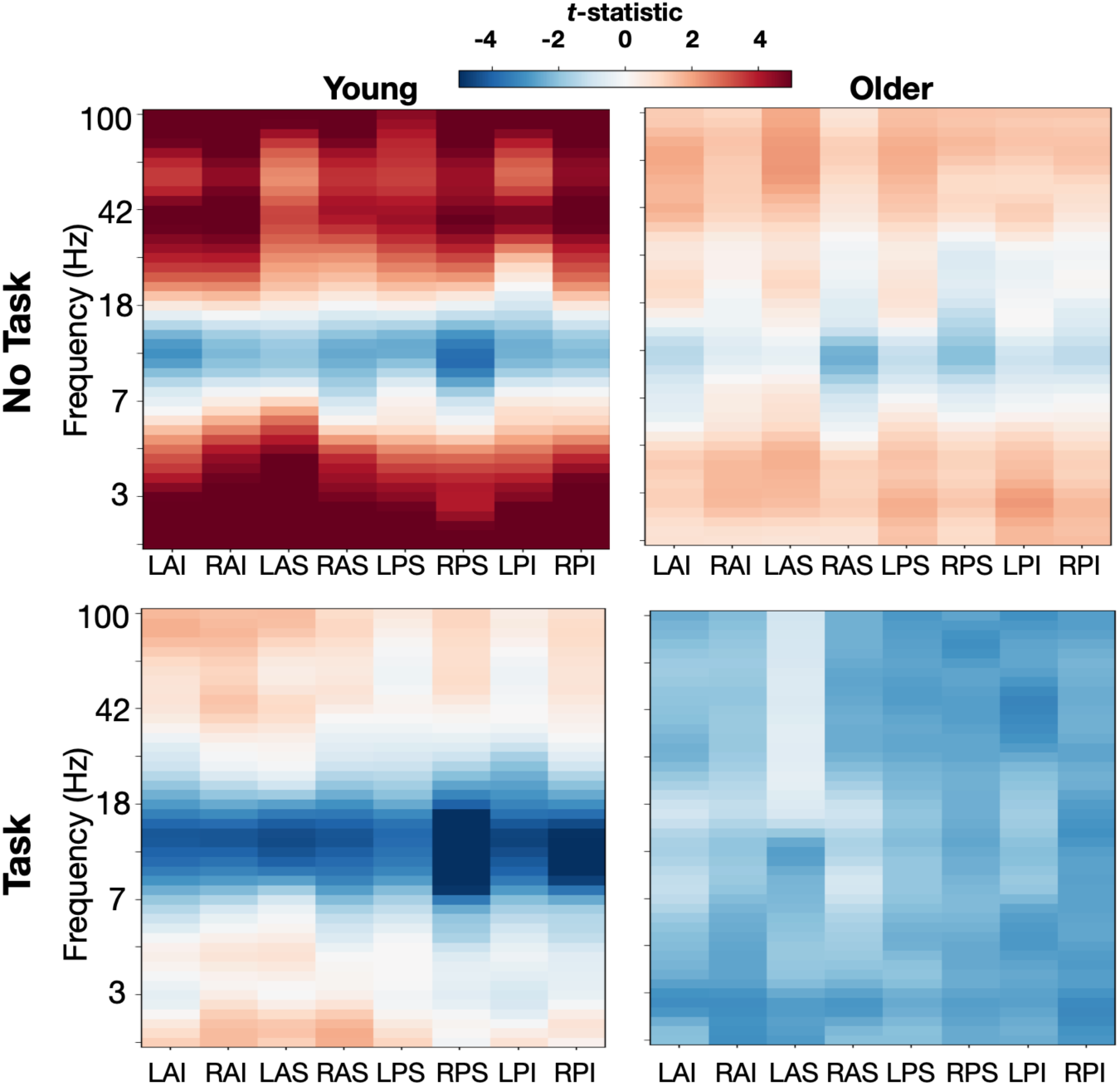
SMEs by List Type. Across-subject t-statistic for recalled - nonrecalled z-power at each computed frequency and ROI. SMEs are computed separately by age group (young/older) and by list condition (no task/task).

We hypothesized that on No Task lists, older adults may demonstrate a reduced gamma SME compared to young adults, which would be consistent with increased taxation of attention during encoding (Broitman and Swallow, 2023; Sederberg et al., 2006; Long and Kahana, 2017). In contrast with this prediction, gamma subsequent memory effects did not vary between age groups in either the Task or No Task lists, age *×* recall status interaction, *F ^′^s <* 1.6*, p^′^s > .*20. However, there was a larger effect of list condition on gamma-band SMEs among older adults than young adults, age *×* task *×* recall status interaction, *F* (1, 205780) = 7.1*, p* = .007. The results therefore suggest that both gamma and alpha activity may be more sensitive to task demands in older than younger adults, and contrast with our observation that older adults’ recall varied *less* across list conditions than that of young adults.

If neural classifiers can predict encoding with similar accuracy among young and older subjects, can they also capture the interacting effects of age group and list condition on memory performance? Figure 5A presents classifier output by age group and list condition. Although we could not evaluate the main effect of age group due to classifier balancing (i.e., classifier outputs average .50 for all subjects), we were still able to model effects of list condition and its interaction with age. We therefore constructed a mixed-effects linear model with classifier output as the dependent variable, fixed effects of age group and list condition, and a random effect of list condition nested within subjects. Across all subjects, classifier output was significantly lower in Task lists than in No Task lists, *F* (1, 203754) = 1299*, p < .*0001. This result is consistent with the main effect of list condition that we also observed in the recall data. Importantly, the difference in classifier output between Task and No Task lists was smaller among the older adult subjects, mirroring the interaction of age group and list condition observed in the recall data, *F* (1, 203754) = 5.3*, p < .*05. These effects demonstrate that EEG spectral activity can capture interacting effects of age and task demands.

**Figure 5.**
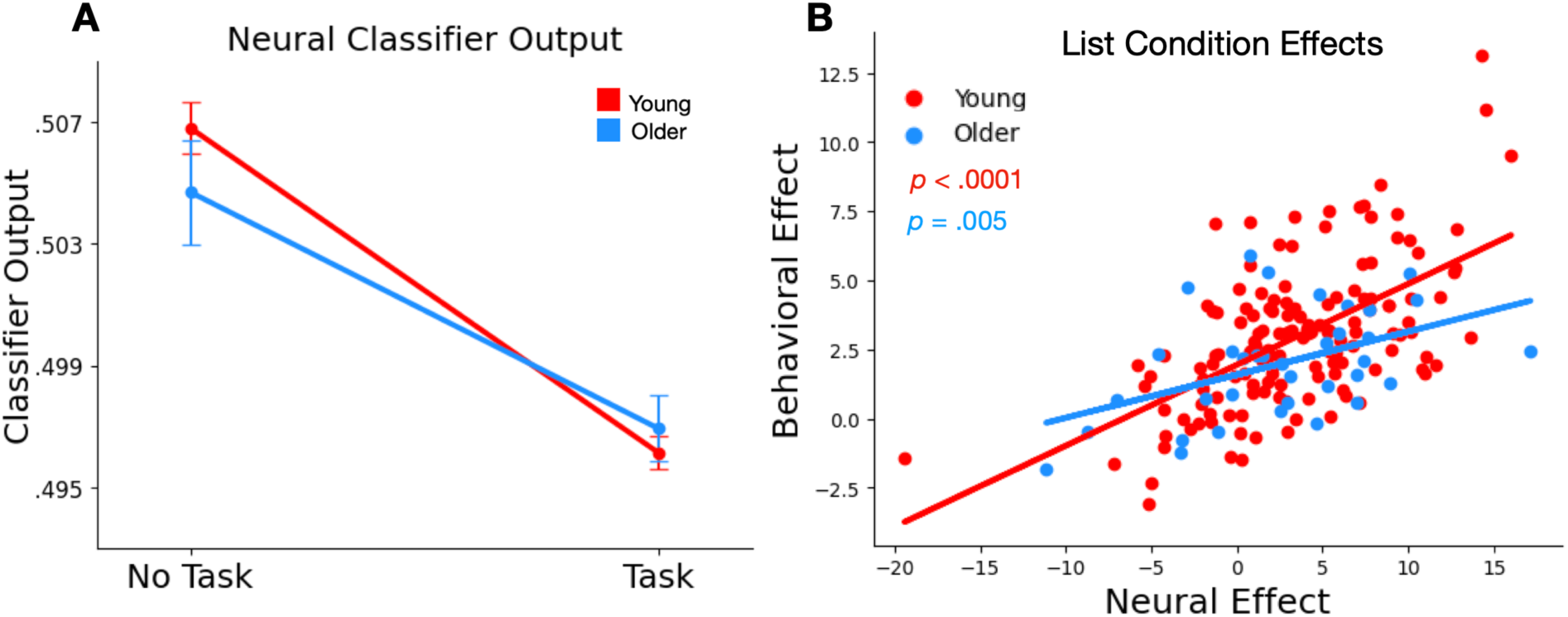
Multivariate classification of EEG data. We trained multivariate logistic models on neural data to predict whether each word would be recalled later. **A.** Average classifier output, plotted separately by age group and list condition. **B.** Correlating Neural and Behavioral Markers of Task Demands. Each subject’s independent samples t-tests on classifier outputs from No Task vs. Task lists, compared with their corresponding t-tests on recall for No Task vs. Task lists. Pearson correlations of the two measures revealed a strong, significant relationship between behavioral and neural effects of list condition in both young adults (r = .58) and older adults (r = .46).

The neural classifiers, which were trained only to predict the binary label of subsequent recall, nonetheless captured the interacting effects of age and list condition on memory at the group level. To determine whether classifiers reflected the effects of task demands at the individual level, we correlated each subject’s behavioral effects of list condition with their corresponding change in neural activity. For each individual subject, we performed independent samples *t*-tests on their recall data between the list conditions, as well as their neural classifier output between list conditions. Figure 5B presents a scatter plot of these *t*-statistics for each subject. We found that behavioral and neural effects of list condition were strongly correlated, suggesting that the classifiers were able to capture the effects of task demands at the individual level. Although this correlation was numerically stronger among the young adult subjects than the older ones (young adults: *r* = .58*, p < .*0001; older adults: *r* = .46*, p* = .005), the correlations did not significantly differ across age groups, *t*(177) = .84*, p* = .39.

## Discussion

In the present study, we investigated whether we can capture the effects of aging and task demands on memory performance using multivariate decomposition of scalp EEG signals. Specifically, we sought to evaluate whether EEG features related to memory encoding varied between older and young adults when they encoded word lists either under full attention or while performing a secondary semantic processing task. Compared to young adults, older adults demonstrated impaired memory performance, but age differences were smaller when subjects performed a secondary task at encoding. Our analyses revealed that older adults exhibit distinct SMEs, characterized by less negativity in the alpha frequencies. Gamma band SMEs reflected changes in task demands, particularly among older adults. Finally, our multivariate EEG classifiers predicted encoding success with similar accuracy in both young and older adults, and reflected the interaction of age and task demands on memory performance.

### Interacting effects of aging and semantic processing on memory encoding

Our free recall data replicated prior findings that both aging and increasing task demands correspond with reduced free recall performance. However, older adults’ memory performance was closer to that of the young adults when they had to perform a secondary task at encoding. This result is surprising, given evidence that older adults are particularly vulnerable to dual-task interference while encoding verbal materials (Park et al., 1989; Kemper et al., 2010). The non-replication of these findings in the current study may be due to the secondary task requiring subjects to make a semantic judgement about each of the to-be-encoded words. This process likely disrupts the rehearsal of early list items, an efficient encoding strategy that young adults frequently use, but which older adults use less frequently (Sanders et al., 1980; Jackson and Schneider, 1982; Ward and Maylor, 2005). A secondary task that selectively impairs rehearsal could therefore be less disruptive for older adults than for young ones. The reduced primacy gradient on Task lists among both age groups, observable in Fig. 2B, supports this explanation.

Another possibility is that the age*×*task interaction on recall reflects reduced cognitive flexibility among older adults. As the experiment switched between Task and No Task lists, younger subjects may have more effectively adopted encoding strategies (e.g., covert rehearsal) that take advantage of the reduced cognitive load from the lack of a secondary task (Mundorf et al., 2022). This would explain the larger recall differences between the age groups in No Task lists, and is consistent with a wealth of literature reporting that cognitive aging accompanies reductions in cognitive flexibility and the ability to switch between tasks (Kramer et al., 1999; Kray and Lindenberger, 2000; Wecker et al., 2005; West and Travers, 2008). The results add nuance to the literature regarding the effects of cognitive aging on the ability to flexibly divide attention across tasks, demonstrating the variability of these effects across task types.

### Gamma SMEs vary with task demands

Our EEG analysis demonstrated that the relationship between gamma power and memory varies with task demands during encoding. Gamma power positively predicted mnemonic success during No Task lists, but *negatively* predicted it on Task lists. This result aligns with Broitman & Swallow’s 2023 finding that the gamma SME was positive when subjects encoded word lists under full attention, but absent when they performed a continuous target detection task during encoding. Considered with other literature indicating gamma oscillations are involved in sustaining attention over time (Clayton et al., 2015; Randazzo et al., 2019; Sederberg et al., 2006), our results support the hypothesis that performing multiple tasks simultaneously disrupts the maintenance of task-oriented states.

Within each list condition, the relationship between gamma power and encoding success remained consistent across age groups. However, a three-way interaction of recall status, age, and list condition indicated that the older adults’ gamma SMEs demonstrated a greater change across list types than those of the young adults. Furthermore, older adults exhibited larger increases in alpha power across list conditions than the young adults did. These results are surprising, because older adults’ recall was more similar across list types than that of the young adults. This finding could indicate that gamma and alpha activity become more sensitive to task demands with age. Supporting this hypothesis is the prior finding that older adults exhibit a steeper gamma primacy gradient during word list encoding, with power declining rapidly across serial positions as they divide their attention to simultaneously process multiple items (Healey and Kahana, 2020).

### Older adults exhibit distinct spectral SMEs

Alpha suppression had weaker associations with mnemonic success in older adults compared to young adults. Given existing evidence that band-averaged alpha power decreases with adult aging (Polich, 1997; Scally et al., 2018; Tröndle et al., 2023), it is unsurprising that alpha’s relationship to memory encoding may diminish with age. Although it remains unclear exactly how aging might reduce the alpha SME, the inability to modulate alpha has been linked with inhibitory deficits in older adults (Vaden et al., 2012; Li and Zhao, 2015; Schmiedt-Fehr et al., 2016). The reduced alpha SME among older adults may therefore relate to increased difficulty suppressing task-irrelevant information at encoding, a factor proposed to underlie age-related memory decline (Hasher and Zacks, 1988; Craik and Byrd, 1982; Healey and Kahana, 2016).

Inhibition deficit frameworks of cognitive aging (Craik and Byrd, 1982; Hasher and Zacks, 1988; Schmitz et al., 2010; Hasher and Campbell, 2020) attribute age-related memory impairment to a reduced ability to suppress the processing of task-irrelevant information. These theories might predict that imposing a secondary task during encoding could simulate cognitive aging effects (Greene et al., 2020). Behaviorally, during Task lists young adults exhibited reductions in overall recall and primacy effects (Fig. 2A), both linked to cognitive aging (Healey and Kahana, 2016; Bruno et al., 2013). However, the *neural correlates* of memory encoding among young adults in the Task lists did not resemble those of older adults in the No Task lists; instead of alpha suppression becoming less predictive of memory encoding, the gamma SME changed from positive to negative. This result indicates that imposing a semantic processing task on young adults does not accurately simulate the effects of cognitive aging on memory encoding. Rather, some of the age-related encoding deficits may be due to additional processes beyond the inhibition of task-irrelevant information. Further work examining the varying effects of task demands on the spectral biomarkers of memory is needed to pursue these possibilities.

### Multivariate classifiers capture interactions of aging and task demands

Multivariate neural classifiers predicted memory with similar accuracy across age groups, and captured interactions of age and task demands in the EEG data. Unlike group-level analyses, EEG classifiers provide individualized models that relate neural activity to mnemonic success at the trial level. We leveraged these models to investigate whether individual differences in the change in recall performance across No Task and Task lists correlated with the degree to which the classifiers’ prediction of encoding success varied across list conditions. Our results indicate that the classifiers not only captured the interacting effects of age and task demands on memory, but also that each subject’s change in memory performance across conditions was strongly correlated with changes in classifier output. These findings suggest that the classification-based model of encoding success incorporates features that reflect both cognitive aging and task demands.

In addition to capturing effects of aging and task demands, mnemonic classifiers are capable of predicting temporal clustering effects (Li et al., 2024) and the effects of neural stimulation on memory encoding (Ezzyat et al., 2018). Future studies may seek to leverage classifiers to study the effects of aging on other cognitive processes related to memory, such as skill acquisition (Broitman et al., 2020) or temporal clustering (Talamonti et al., 2020).

### Study limitations

A potential confound concerns the fact that Task lists differed from No Task lists by requiring subjects to perform a physical button press during each word presentation, introducing the possibility for motor artifacts which could influence EEG activity. However, because all trials in Task lists required a button press (irrespective of whether they were recalled or nonrecalled), the SME analysis should have regressed out any resulting motor artifacts. Furthermore, motor effects from a button press would likely appear in scalp sites contralateral to the hand performing the action (Quandt et al., 2012; Deiber et al., 2012). Although we did not instruct subjects to use one hand over the other, all subjects in the current study were right-handed and performed the task using adjacent keys on the right side of the keyboard, so they would have likely used their dominant hand to make responses. However, the effect of list condition on the gamma SMEs was observed diffusely across bilateral scalp sites (see Fig. 4). Motor artifacts are thus unlikely to explain the differential gamma SMEs observed across list conditions.

An additional limitation concerns the fact that we did not explicitly design the present study to test the effects of dual-task interference on memory performance across age groups. Rather, we intended to investigate the effects of semantic processing and experimental task context on the behavioral and electrophysiological features of free recall (Long and Kahana, 2017; Lohnas et al., 2023; Polyn et al., 2012). We therefore cannot broadly generalize the results to all dual-task encoding paradigms, nor can we conclusively determine whether any effects of list type were due to dual-task interference or added semantic processing. Additional investigations are therefore necessary to determine the specific effects of dual-task interference on the neural biomarkers of encoding, and whether such effects vary across the adult life span.

### Conclusions

Both cognitive aging and increasing task demands impair memory encoding. We found that EEG alpha activity reflected the effects of aging on memory, and both gamma and alpha oscillations captured effects of task demands. Differences in recall between the age groups were smaller when participants performed semantic judgments on each word during encoding. Classifiers trained on the neural data successfully predicted memory encoding at the trial level in both young and older adults and captured interactions of age and task demands.

## References

1. Babiloni, C., Binetti, G., Cassarino, A., Dal Forno, G., Del Percio, C., Ferreri, F., Ferri, R., Frisoni, G., Galderisi, S., Hirata, K., et al. (2006). Sources of cortical rhythms in adults during physiological aging: a multicentric eeg study. Human brain mapping, 27(2):162–172.

2. Bastiaansen, M., Mazaheri, A., and Jensen, O. (2011). Beyond erps: oscillatory neuronal. Oxf. Handb. Event Relat. Potential Compon, pages 31–50.

3. Bates, D., Maechler, M., Bolker, B., Walker, S., Christensen, R. H. B., Singmann, H., Dai, B., Grothendieck, G., Green, P., and Bolker, M. B. (2015). Package ?lme4? convergence, 12(1):2.

4. Broitman, A. W., Kahana, M. J., and Healey, M. K. (2020). Modeling retest effects in a longitudinal measurement burst study of memory. Computational brain & behavior, 3(2):200–207.

5. Broitman, A. W. and Swallow, K. M. (2023). Temporal attention modulates eeg spectral correlates of successful memory encoding. bioRxiv, pages 2023–04.

6. Bruno, D., Reiss, P. T., Petkova, E., Sidtis, J. J., and Pomara, N. (2013). Decreased recall of primacy words predicts cognitive decline. Archives of clinical neuropsychology, 28(2):95–103.

7. Cabeza, R., Daselaar, S. M., Dolcos, F., Prince, S. E., Budde, M., and Nyberg, L. (2004). Task-independent and task-specific age effects on brain activity during working memory, visual attention and episodic retrieval. Cerebral cortex, 14(4):364–375.

8. Clayton, M. S., Yeung, N., and Kadosh, R. C. (2015). The roles of cortical oscillations in sustained attention. Trends in cognitive sciences, 19(4):188–195.

9. Craik, F. I. M. and Byrd, M. (1982). Aging and cognitive deficits: The role of attentional resources. In Craik, F. I. M. and Trehub, S., editors, Aging & Cognitive Processes: Advances in the Study of Communication and Affect, volume 8, pages 191–211. Plenum Press, New York.

10. Daselaar, S. M., Fleck, M. S., Dobbins, I. G., Madden, D. J., and Cabeza, R. (2006). Effects of healthy aging on hippocampal and rhinal memory functions: an event-related fmri study. Cerebral cortex, 16(12):1771–1782.

11. Deiber, M.-P., Sallard, E., Ludwig, C., Ghezzi, C., Barral, J., and Ibañez, V. (2012). Eeg alpha activity reflects motor preparation rather than the mode of action selection. Frontiers in integrative neuroscience, 6:59.

12. Dennis, N. A., Kim, H., and Cabeza, R. (2007). Effects of aging on true and false memory formation: An fmri study. Neuropsychologia, 45(14):3157–3166.

13. Ezzyat, Y., Wanda, P., Levy, D., Kadel, A., Aka, A., Pedisich, I., Sperling, M., Sharan, A., Lega, B., Burks, A., Gross, R., Inman, C., Jobst, B., Gorenstein, M., Davis, K., et al. (2018). Closed-loop stimulation of temporal cortex rescues functional networks and improves memory. Nature Communications, 9(365).

14. Finnigan, S., O’Connell, R. G., Cummins, T. D., Broughton, M., and Robertson, I. H. (2011). Erp measures indicate both attention and working memory encoding decrements in aging. Psychophysiology, 48(5):601–611.

15. Friedman, D. (2013). The cognitive aging of episodic memory: a view based on the event-related brain potential. Frontiers in behavioral neuroscience, 7:111.

16. Greene, N. R., Naveh-Benjamin, M., and Cowan, N. (2020). Adult age differences in working memory capacity: Spared central storage but deficits in ability to maximize peripheral storage. Psychology and Aging, 35(6):866.

17. Hasher, L. and Campbell, K. L. (2020). Inhibitory theory: Assumptions, findings, and relevance to interventions.

18. Hasher, L. and Zacks, R. T. (1988). Working memory, comprehension, and aging: A review and a new view. In Bower, G. H., editor, The psychology of learning and motivation: Advances in research and theory, pages 193–225. Academic Press, San Diego.

19. Haufe, S., Meinecke, F., Görgen, K., Dähne, S., Haynes, J.-D., Blankertz, B., and Bießmann, F. (2014). On the interpretation of weight vectors of linear models in multivariate neuroimaging. NeuroImage, 87:96–110.

20. Healey, M. K. and Kahana, M. J. (2016). A four–component model of age–related memory change. Psychological Review, 123:23–69.

21. Healey, M. K. and Kahana, M. J. (2020). Age-related differences in the temporal dynamics of spectral power during memory encoding. Plos one, 15(1):e0227274.

22. Jackson, D. and Schneider, H. (1982). Age differences in organization and recall: An analysis of rehearsal processes. Psychological Reports, 50(3):919–924.

23. Jacobs, J., Hwang, G., Curran, T., and Kahana, M. J. (2006). EEG oscillations and recognition memory: Theta correlates of memory retrieval and decision making. NeuroImage, 15(2):978–87.

24. Kahana, M. J., Lohnas, L. J., Healey, K., Aka, A., Broitman, A., Crutchley, E., Crutchley, P., Alm, K. H., Katerman, B. S., Miller, N. E., et al. (2022). The penn electrophysiology of encoding and retrieval study.

25. Karrasch, M., Laine, M., Rapinoja, P., and Krause, C. M. (2004). Effects of normal aging on event-related desynchronization/synchronization during a memory task in humans. Neuroscience letters, 366(1):18–23.

26. Katerman, B. S., Li, Y., Pazdera, J. K., Keane, C., and Kahana, M. J. (2022). Eeg biomarkers of free recall. NeuroImage, 246:118748.

27. Kemper, S., Schmalzried, R., Hoffman, L., and Herman, R. (2010). Aging and the vulnerability of speech to dual task demands. Psychology and aging, 25(4):949.

28. Kim, H. (2011). Neural activity that predicts subsequent memory and forgetting: a meta-analysis of 74 fMRI studies. NeuroImage, 54(3):2446–2461.

29. Kragel, J. E., Ezzyat, Y., Sperling, M. R., Gorniak, R., Worrell, G. A., Berry, B. M., Inman, C., Lin, J.-J., Davis, K. A., Das, S. R., Stein, J. M., Jobst, B. C., Zaghloul, K. A., Sheth, S. A., Rizzuto, D. S., and Kahana, M. J. (2017). Similar patterns of neural activity predict memory function during encoding and retrieval. NeuroImage, 155:60–71.

30. Kramer, A. F., Hahn, S., and Gopher, D. (1999). Task coordination and aging: Explorations of executive control processes in the task switching paradigm. Acta psychologica, 101(2-3):339–378.

31. Kray, J. and Lindenberger, U. (2000). Adult age differences in task switching. Psychology and aging, 15(1):126.

32. Kukolja, J., Göreci, D. Y., Onur, O. A., Riedl, V., and Fink, G. R. (2016). Resting-state fmri evidence for early episodic memory consolidation: effects of age. Neurobiology of aging, 45:197–211.

33. Lacombe, J., Jolicoeur, P., Grimault, S., Pineault, J., and Joubert, S. (2015). Neural changes associated with semantic processing in healthy aging despite intact behavioral performance. Brain and language, 149:118–127.

34. Li, J., Morcom, A. M., and Rugg, M. D. (2004). The effects of age on the neural correlates of successful episodic retrieval: an erp study. *Cognitive, Affective*, & Behavioral Neuroscience, 4(3):279–293.

35. Li, L. and Zhao, D. (2015). Age-related inter-region eeg coupling changes during the control of bottom–up and top–down attention. Frontiers in aging neuroscience, 7:223.

36. Li, Y., Pazdera, J. K., and Kahana, M. J. (2024). EEG decoders track memory dynamics. Nature Communications, 15(1):2981.

37. Lohnas, L. J., Healey, M. K., and Davachi, L. (2023). Neural temporal context reinstatement of event structure during memory recall. Journal of Experimental Psychology: General, 152(7):1840.

38. Long, N. M., Burke, J. F., and Kahana, M. J. (2014). Subsequent memory effect in intracranial and scalp EEG. NeuroImage, 84:488–494.

39. Long, N. M. and Kahana, M. J. (2017). Modulation of task demands suggests that semantic processing interferes with the formation of episodic associations. *Journal of Experimental Psychology: Learning*, Memory, and Cognition, 43(2):167.

40. Mayr, U. and Kliegl, R. (2000). Complex semantic processing in old age: Does it stay or does it go? Psychology and aging, 15(1):29.

41. McBride, J. C., Zhao, X., Munro, N. B., Smith, C. D., Jicha, G. A., Hively, L., Broster, L. S., Schmitt, F. A., Kryscio, R. J., and Jiang, Y. (2014). Spectral and complexity analysis of scalp eeg characteristics for mild cognitive impairment and early alzheimer’s disease. Computer methods and programs in biomedicine, 114(2):153–163.

42. Mundorf, A. M., Uitvlugt, M. G., and Healey, M. K. (2022). Does depth of processing affect temporal contiguity? Psychonomic Bulletin & Review, 29(6):2229–2239.

43. Murty, D. V., Manikandan, K., Kumar, W. S., Ramesh, R. G., Purokayastha, S., Javali, M., Rao, N. P., and Ray, S. (2020). Gamma oscillations weaken with age in healthy elderly in human eeg. NeuroImage, 215:116826.

44. Nelson, D. L., McEvoy, C. L., and Schreiber, T. A. (2004). The University of South Florida free association, rhyme, and word fragment norms. *Behavior Research Methods*, Instruments and Computers, 36(3):402–407.

45. Nilsson, L.-G. (2003). Memory function in normal aging. Acta Neurologica Scandinavica, 107:7–13.

46. Nolan, H., Whelan, R., and Reilly, R. (2010). Faster: fully automated statistical thresholding for eeg artifact rejection. Journal of neuroscience methods, 192(1):152–162.

47. Paller, K. A., Kutas, M., and Mayes, A. R. (1987). Neural correlates of encoding in an incidental learning paradigm. 67:360–371.

48. Park, D. C., Smith, A. D., Dudley, W. N., and Lafronza, V. N. (1989). Effects of age and a divided attention task presented during encoding and retrieval on memory. Journal of Experimental Psychology: Learning, Memory, and Cognition, 15(6):1185.

49. Phan, T. D., Wachter, J. A., Solomon, E. A., and Kahana, M. J. (2019). Multivariate stochastic volatility modeling of neural data. Elife, 8:e42950.

50. Polich, J. (1997). On the relationship between eeg and p300: individual differences, aging, and ultradian rhythms. International journal of psychophysiology, 26(1-3):299–317.

51. Polyn, S. M., Kragel, J. E., Morton, N. W., McCluey, J. D., and Cohen, Z. D. (2012). The neural dynamics of task context in free recall. Neuropsychologia, 50(4):447–457.

52. Quandt, F., Reichert, C., Hinrichs, H., Heinze, H.-J., Knight, R. T., and Rieger, J. W. (2012). Single trial discrimination of individual finger movements on one hand: a combined meg and eeg study. NeuroImage, 59(4):3316–3324.

53. Randazzo, M. J., Ezzyat, Y., and Kahana, M. J. (2019). Spectral tilt underlies mathematical problem solving. bioRxiv, page 601880.

54. Rhodes, S., Greene, N. R., and Naveh-Benjamin, M. (2019). Age-related differences in recall and recognition: A meta-analysis. Psychonomic Bulletin & Review, 26(5):1529–1547.

55. Salthouse, T. A. (2016). Continuity of cognitive change across adulthood. Psychonomic bulletin & review, 23:932–939.

56. Sanders, R. E., Murphy, M. D., Schmitt, F. A., and Walsh, K. K. (1980). Age differences in free recall rehearsal strategies. Journal of Gerontology, 35(4):550–558.

57. Scally, B., Burke, M. R., Bunce, D., and Delvenne, J.-F. (2018). Resting-state eeg power and connectivity are associated with alpha peak frequency slowing in healthy aging. Neurobiology of aging, 71:149–155.

58. Schmiedt-Fehr, C., Mathes, B., Kedilaya, S., Krauss, J., and Basar-Eroglu, C. (2016). Aging differentially affects alpha and beta sensorimotor rhythms in a go/nogo task. Clinical Neurophysiology, 127(10):3234–3242.

59. Schmitz, T. W., Cheng, F. H., and De Rosa, E. (2010). Failing to ignore: paradoxical neural effects of perceptual load on early attentional selection in normal aging. Journal of Neuroscience, 30(44):14750–14758.

60. Sederberg, P. B., Gauthier, L. V., Terushkin, V., Miller, J. F., Barnathan, J. A., and Kahana, M. J. (2006). Oscillatory correlates of the primacy effect in episodic memory. NeuroImage, 32(3):1422–1431.

61. Steyvers, M., Shiffrin, R. M., and Nelson, D. L. (2004). Word association spaces for predicting semantic similarity effects in episodic memory. In Healy, A. F., editor, Cognitive Psychology and its Applications: Festschrift in Honor of Lyle Bourne, Walter Kintsch, and Thomas Landauer. American Psychological Association, Washington, DC.

62. Strunk, J. and Duarte, A. (2019). Prestimulus and poststimulus oscillatory activity predicts successful episodic encoding for both young and older adults. Neurobiology of aging, 77:1–12.

63. Talamonti, D., Montgomery, C. A., Clark, D. P., and Bruno, D. (2020). Age-related prefrontal cortex activation in associative memory: An fnirs pilot study. NeuroImage, 222:117223.

64. Tröndle, M., Popov, T., Pedroni, A., Pfeiffer, C., Barańczuk-Turska, Z., and Langer, N. (2023). Decomposing age effects in eeg alpha power. Cortex, 161:116–144.

65. Vaden, R. J., Hutcheson, N. L., McCollum, L. A., Kentros, J., and Visscher, K. M. (2012). Older adults, unlike younger adults, do not modulate alpha power to suppress irrelevant information. NeuroImage, 63(3):1127–1133.

66. Verhaeghen, P. (2003). Aging and vocabulary scores: A meta-analysis. Psychology and Aging, 18(2):332–339.

67. Voytek, B., Kramer, M. A., Case, J., Lepage, K. Q., Tempesta, Z. R., Knight, R. T., and Gazzaley, A. (2015). Age-related changes in 1/f neural electrophysiological noise. The Journal of Neuroscience, 35(38):13257–13265.

68. Ward, G. and Maylor, E. (2005). Age-related deficits in free recall: The role of rehearsal. Quarterly Journal Of Experimental Psychology, 58A(1):98–119.

69. Wecker, N. S., Kramer, J. H., Hallam, B. J., and Delis, D. C. (2005). Mental flexibility: age effects on switching. Neuropsychology, 19(3):345.

70. Weidemann, C. T. and Kahana, M. J. (2021). Neural measures of subsequent memory reflect endogenous variability in cognitive function. *Journal of Experimental Psychology: Learning*, Memory, and Cognition, 47(4):641.

71. Weidemann, C. T., Mollison, M. V., and Kahana, M. J. (2009). Electrophysiological correlates of high-level perception during spatial navigation. Psychonomic Bulletin & Review, 16(2):313–319.

72. West, R. and Travers, S. (2008). Differential effects of aging on processes underlying task switching. Brain and Cognition, 68(1):67–80.

